# Genomic characterization and evolutionary path of human satellite 1B in primates highlight a dynamic relationship with transposable elements

**DOI:** 10.64898/2026.11.05.760144

**Authors:** Carlos Vila-Verde, Alessia Daponte, Daniel Eleutério, Raquel Chaves, Claudia Rita Catacchio, Sandra Louzada

## Abstract

Human Satellite 1B (HSat1B) remains one of the least characterized classical human satellite DNAs, despite its massive abundance on the human Y chromosome. Its tripartite ∼2.5 kb monomer - comprising an *AluY* fragment, a GC-rich unit (here named H1B) and an AT-rich block - points to an evolutionary origin tightly linked to transposable elements (TEs).

Here, we present the first comprehensive comparative genomic, phylogenetic, and transcriptional characterization of HSat1B across 10 primate species using Telomere-to-Telomere (T2T) and chromosome-scale long-read assemblies, combined with DNA-FISH, RNA-FISH and long-read transcriptomic analyses. We demonstrate that while canonical tandem HSat1B arrays are restricted to Homininae (human, chimpanzee, bonobo, and gorilla), the core H1B unit and its structural variants (H1Bsv1 and H1Bsv2) occur across all 10 primate genomes, including strepsirrhines (lemur). This indicates that the H1B sequence predates the assembly of the composite HSat1B monomer by at least 60 million years. H1B variants consistently map within TE-rich genomic contexts, supporting a stepwise, TE-mediated model of satellite diversification and mobilization.

Within Homininae, humans exhibit a dramatic lineage-specific expansion driven by Yq12 amplification (∼14.2 Mb; 92% of total genomic content), whereas non-human great apes carry smaller, highly polymorphic autosomal and Y-linked arrays. Finally, we show that H1B-containing sequences are transcriptionally active in human and gorilla nuclei via RNA-FISH and characterize heterogeneous H1B/*Alu* chimeric non-coding transcripts in the human male germline. Collectively, our findings uncover the birth and Homininae-specific expansion of a composite satellite DNA seeded by an ancient primate sequence module and sculpted by transposable elements.

## Introduction

The repetitive fraction of eukaryotic genomes, long dismissed as non-functional DNA (Doolittle, 2013; Palazzo & Gregory, 2014), has emerged as a functional and evolutionary component of genome architecture. Among repetitive sequences, satellite DNAs (satDNAs) are characterized by tandemly arranged repeat units organized in large arrays, predominantly in heterochromatic regions, where they contribute to centromere function, chromosome structure and genome stability (Garrido-Ramos, 2017). Moreover, because satDNA sequences evolve rapidly, they are increasingly recognized as primary drivers of genomic and karyotypic diversification (Louzada et al., 2020). In primates, satDNA families show lineage-specificity in their sequence composition, chromosomal distribution and array sizes (Ahmad et al., 2020). The recent completion of Telomere-to-Telomere (T2T) genome assemblies for human and multiple non-human primates (Nurk et al., 2022; Rhie et al., 2023; Yoo et al., 2025) has opened many opportunities to study these previously inaccessible genomic regions at single-nucleotide resolution.

The satellite DNA sequence currently designated as human satellite 1B (HSat1B) was originally identified as male-specific and annotated as HSATI (Cooke et al., 1982; Manuelidis, 1976) and described as a component of a ∼2.5 kb tripartite monomer consisting of an AT-rich region, an Alu element and a GC-rich unit (Frommer et al., 1984). While early studies localized HSat1B to the human Y and acrocentric chromosomes, it was later identified across other primate genomes (Babcock et al., 2007). This satellite was only recently renamed HSat1B to distinguish it from HSat1A, another distinct AT-rich satellite family and it has been characterized together with other human satellites in a recent study (Altemose et al., 2022).

The presence of an Alu element within the HSat1B monomer is noteworthy. Alu fragments are primate-specific transposable elements (TEs) that have contributed extensively to genome remodeling (Sundaram et al., 2014). The association between TEs and satDNA arrays has been documented across lineages and is proposed as a pathway through which new satellite families emerge and diversify (Šatović-Vukšić & Plohl, 2023).

Satellite DNAs relevance in the genomes has been related also to their transcription into satellite non-coding RNAs (satncRNAs) with roles in chromatin regulation, centromere maintenance and nuclear stress responses (Hoyt et al., 2022; Ninomiya et al., 2023), but the transcriptional activity of HSat1B across primate species has not yet been studied. Overall, the tripartite organization of HSat1B, its enrichment on the human Y chromosome and its association with TE-derived sequence blocks, makes HSat1B an interesting sequence for investigating how satellite DNA can emerge and diversify in primate genomes.

Here, we present a comprehensive comparative genomic analysis of HSat1B across 10 primate species, from lemurs to humans, integrating long read genome assemblies with DNA-FISH, RNA-FISH and phylogenetic inference. HSat1B transcription analysis was complemented with available transcriptomic data revealing HSat1B transcript’s structure. Our data allowed us to explore the evolutionary pathway of this satellite across primate evolution being its origin and expansion tightly associated to TE elements.

## Results

### Human genome contains HSat1B satellite and HSat1B-related sequences

Our analysis began by gathering all satellite DNA annotations from the CenSat and RepeatMasker tracks in the UCSC Genome Browser (https://www.genome.ucsc.edu) for the T2T-CHM13v2.0 assembly. This revealed the existence of HSat1B spanning 66 different locations on the human genome distributed across 6 chromosomes (Table 1, Supplementary Table S1a).

**Table 1.** Genomic distribution, number of arrays and number of monomer found in each human chromosome from T2TCHM13v2.0.

|  | **HSat1B** |  |  |
|---|---|---|---|
|  | **Amount (bp)** | **# Arrays** | **# Monomer** |
| chr13 | 97167 | 4 | 38.9 |
| chr14 | 93127 | 2 | 37.3 |
| chr15 | 444131 | 1 | 177.7 |
| chr21 | 66373 | 3 | 26.5 |
| chr22 | 460281 | 10 | 184.1 |
| chrY | 14206138 | 46 | 5682.5 |

Each monomer consisted of an AluY fragment (∼291 bp), a GC-rich unit, here designated as H1B (∼566 bp) and an AT-rich block (∼1,553 bp), comprising approximately 2.5 Kb (Figure 1A, Supplementary Table S1a). HSat1B was organized in tandem arrays ranging from 3 to 474 monomers per array, with the majority of HSat1B content located on chromosome Yq12 (14.2 Mb) and the remaining arrays distributed across the pericentromeric regions of acrocentric chromosomes 13, 14, 15, 21 and 22 (1.2 Mb in total) (Figure 1D, Table 1, Supplementary Table S1a). HSat1B arrays contiguity was estimated based on the number of locations in each chromosome compared to the total number of monomers contained by each array. Among the acrocentric chromosomes, chromosome 22 was the one with the highest number of arrays (10) contrasting to chromosome 15 which showed only one localization. Despite this difference in number of arrays, they contain a very similar monomer number, 178 (chr15) and 168 (ch22), revealing the highest contiguity of chromosome 15 among all the acrocentric. Chromosome Y showed the highest number of monomers, distributed in 46 different locations.

**Figure 1.**
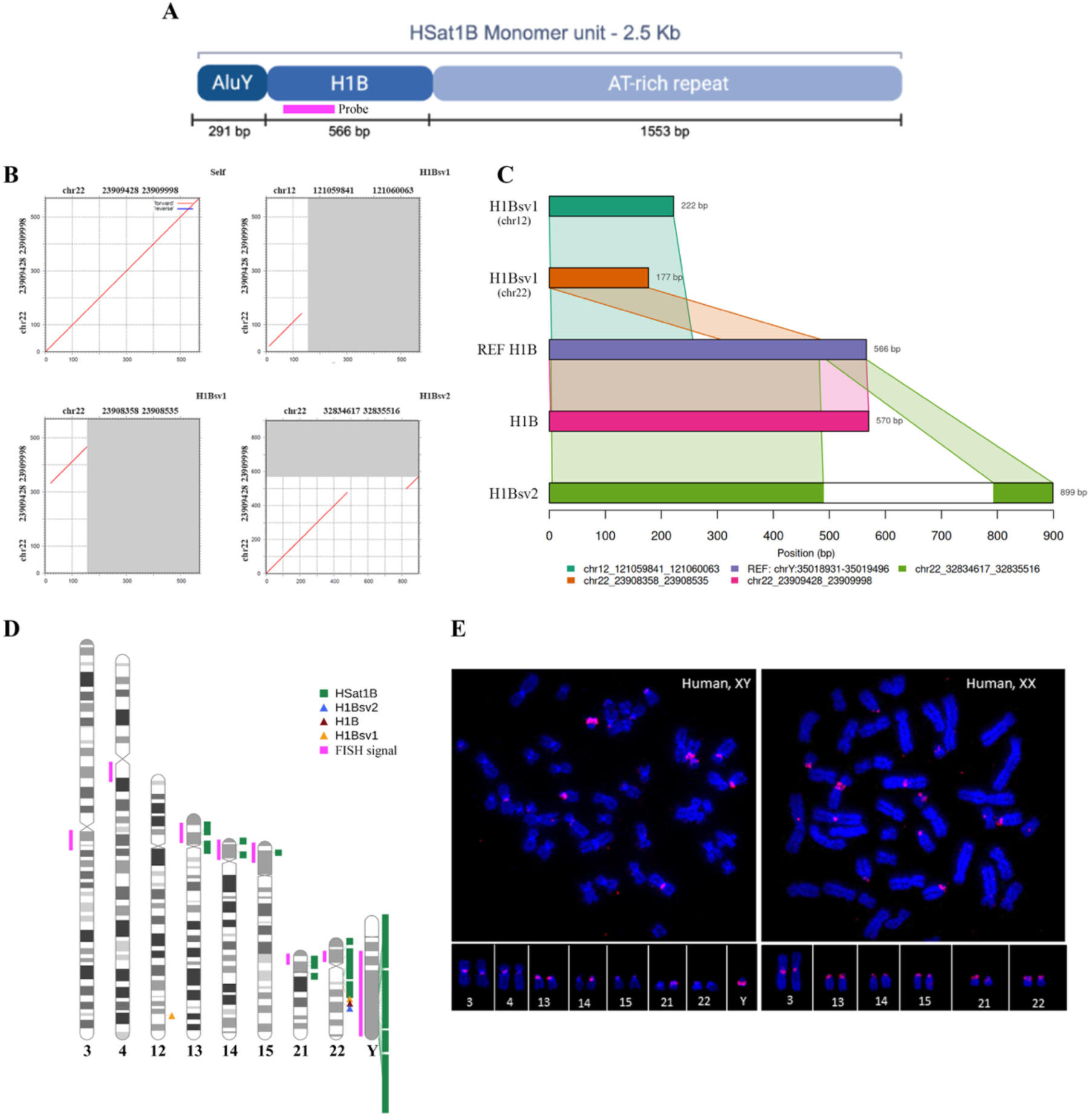
Structural, comparative, and cytogenetic characterization of HSat1B and H1B variants. **(A)** Schematic representation of the canonical HSat1B genomic monomer (∼2.5 kb), organized into three segments: an *AluY* fragment (291 bp), the GC-rich H1B unit (566 bp, annotated as HSATI in Dfam), and an AT-rich repeat region (1,553 bp) in the T2T-CHM13v2.0 assembly. The pink bar marks the position of the fluorescence *in situ* hybridization (FISH) probe targeting the H1B unit. **(B)** Pairwise dotplot matrices generated with the MAFFT web server comparing the reference H1B unit against structural variants identified across the T2T-CHM13v2.0 assembly. The top-left panel shows the self-alignment of the reference H1B locus (chr22:23,909,428–23,909,998), while the remaining panels compare this reference against H1Bsv1 (chr12 and chr22) and H1Bsv2 (chr22) loci (genomic coordinates provided in Supplementary Tables S1a–c). **(C)** Sequence alignment and structural comparison of the H1B unit and its structural variants (H1Bsv1 on chr12 and chr22; H1Bsv2 on chr22) against the annotated chromosome Y H1B reference (chrY:35,018,931–35,019,496). **(D)** RIdeogram visualization mapping all genomic locations of canonical HSat1B arrays (green squares), dispersed H1B units (red triangles), H1Bsv2 (blue triangles), and H1Bsv1 (yellow triangles) across the human karyotype, alongside experimental DNA-FISH hybridization signals (pink bars). **(E)** Representative DNA-FISH metaphase spreads and corresponding karyograms showing hybridization signals obtained with the H1B-targeting probe in human male (HDFn, XY) and female (GM12878, XX) cell lines (chromosomes counterstained with DAPI, blue; probe signal in red). Strong signals validate the *in silico*-predicted arrays on chromosome Yq12 and the pericentromeric regions of acrocentric chromosomes (13, 14, 15, 21, and 22), while additional signals detected on chromosome 3 (both cell lines) and chromosome 4 (male cell line) where no canonical HSat1B arrays are annotated in T2T-CHM13v2.0, likely reflect dispersed H1B-related sequences or inter-individual copy-number variation.

From the HSat1B arrays detected on chromosome 22, some have a very short AT-rich block (∼60-137 bp) when compared to the canonical HSat1B monomer (∼1,500 bp), observed that in these locations the CenSat annotation track does not identify these regions as HSat1B.

In addition to tandemly arranged HSat1B monomers, HSat1B-related sequences were also detected as dispersed single-copy or low-copy elements across the human genome. These sequences correspond to the GC-rich H1B component of the HSat1B monomer, indicating that H1B is present both within composite HSat1B arrays and independently in the genome. Two structural H1B variants, H1Bsv1 and H1Bsv2, were identified. H1Bsv1 is characterized by a shorter H1B sequence, whereas H1Bsv2 contains an Alu-derived insertion within the H1B unit (Figure 1B, 1C). H1Bsv1 was mapped to chr12:121,059,841–121,060,063 and chr22:23,908,358–23,908,535, positioned within ∼1.1 kb of the canonical H1B locus (chr22:23,909,428–23,909,998), while H1Bsv2 was mapped to chr22:32,834,617– 32,835,516 (Figure 1B, 1C; Supplementary Tables S1b and S1c). The flanking regions of HSat1B and H1B sequences were enriched in Alu and LINE-1 elements, as revealed by RepeatMasker annotations (data not shown).

Phylogenetic analyses of H1B units identified within HSat1B arrays (99% identity threshold) placed the majority of chromosome Y-derived H1B copies in two clusters, whereas H1B copies from the acrocentric chromosomes showed a broader distribution across clusters. H1B-Y has therefore undergone recent expansion and homogenization (Supplementary Figure 1), while H1B sequences from different acrocentric chromosomes are frequently indistinguishable at the sequence level, consistent with recent or ongoing inter-chromosomal sequence exchange.

### Physical mapping of HSat1B by DNA-FISH validates and complements *in silico* data

Physical mapping on human chromosomes, was performed using an HSat1B-specific probe prepared from a cloned PCR fragment using specific primers for H1B sequence. The fragment used for the probe was restricted to part of the HSat1B monomer, given it includes Alu and AT simple repeats, that could cross-hybridize to other locations in the genome. DNA-FISH experiments were performed in metaphase spreads obtained from human male and female cell lines, to verify the presence of differential location and abundance between them. Strong hybridization signals were detected on chromosome Y and, at lower intensity, on the pericentromeric regions of acrocentric chromosomes, in accordance with the *in silico* mapping results (Figure 1D and 1E). Interestingly, hybridization signals were also detected on chromosome 3 in both cell lines and on chromosome 4 in the male cell line.

### HSat1B and H1B-related sequences are present in the genome of non-human primates

Following the same methodological approach applied to the human genome, we investigated the presence of HSat1B and HSat1B-related sequences in chromosome-scale assemblies from nine non-human primate species. Assembly names, accession numbers and haplotype representation are provided in Supplementary Table S2.

The complete HSat1B composite monomer was detected exclusively in *Homininae* species (chimpanzee, bonobo and gorilla). In bonobo and chimpanzee, HSat1B was identified on seven autosomes (chromosomes 6, 14, 15, 16, 17, 22 and 23), with chromosome 22 detected only in the bonobo paternal haplotype. Most occurrences were in pericentromeric regions of chromosomes orthologous to the human acrocentric chromosomes (Figure 2). Compared with humans, HSat1B arrays in bonobo and chimpanzee were substantially smaller, ranging from 4 to 28 monomers, with the largest arrays observed on chromosomes 22 and 23 (Supplementary Tables S3a and S3b).

**Figure 2.**
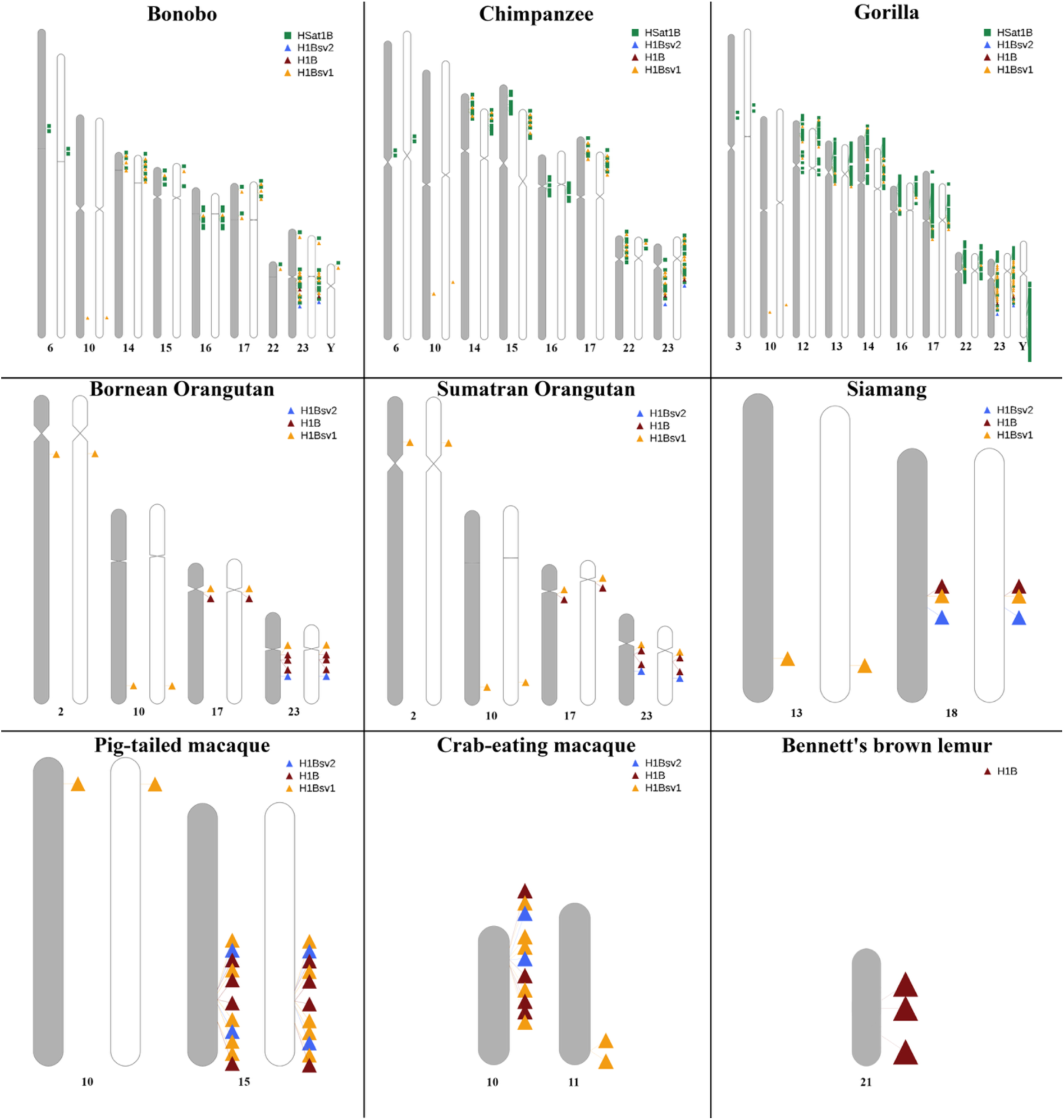
Cross-species chromosomal distribution of HSat1B and H1B sequence variants across non-human primates. RIdeogram visualization mapping the genomic locations of canonical HSat1B arrays (green squares), the H1B unit (red triangles), H1Bsv2 (blue triangles), and H1Bsv1 (yellow triangles) across the karyotypes of nine non-human primate species: bonobo (*Pan paniscus*), chimpanzee (*Pan troglodytes*), gorilla (*Gorilla gorilla*), Bornean orangutan (*Pongo pygmaeus*), Sumatran orangutan (*Pongo abelii*), siamang (*Symphalangus syndactylus*), pig-tailed macaque (*Macaca nemestrina*), crab-eating macaque (*Macaca fascicularis*), and Bennett’s brown lemur (*Eulemur rufifrons*). Color coding matches Figure 1D for direct cross-species comparison. Genome assembly accessions and haplotype details are provided in Supplementary Table S2.

Similarly, in gorilla, HSat1B monomers were identified predominantly in the pericentromeric regions of eight autosomes (chromosomes 3, 12, 13, 14, 16, 17, 22, and 23) and on the Y chromosome (Figure 2), with array sizes varying between 4 and 19 monomers and the most extensive arrays located on chromosome 23 (Supplementary Table S3c). As observed for human chromosome 22, a subset of arrays in bonobo, chimpanzee and gorilla (classified as HSat1B arrays with truncated AT-rich blocks in Supplementary Tables S3a–c) contained a truncated AT-rich repeat compared to the canonical monomer and was therefore missed by automated satellite-annotation pipelines. Nevertheless, BLAST sequence identity and structural continuity with flanking monomers allowed their unambiguous classification as HSat1B. Comparison between maternal and paternal haplotypes revealed high overall concordance in chromosomal distribution, with the notable exception of the absence of HSat1B on the bonobo maternal chromosome 22.

HSat1B-related sequences, including H1B and its structural variants H1Bsv1 and H1Bsv2 - were detected across all nine non-human primate genomes, predominantly as dispersed single- or low-copy elements outside canonical tandem HSat1B arrays (Supplementary Table S3a-i). As depicted in Figure 2, for chimpanzee and bonobo, H1B and H1Bsv2 are restricted to chromosome 23 in both species, while H1Bsv1 shows a more spread localization spanning chromosomes 10, 14, 17, 22, 23 and Y (only for bonobo). Apart from chromosome 10, H1Bsv1 sequences are within the same region of the satellite HSat1B. A similar pattern can be observed in gorilla genome, with H1B and H1Bsv2 restricted to chromosome 23 and H1Bsv1 being in several chromosomes – chr. 10, 12, 13, 14, 16, 17, 22 and 23. Notably, in both orangutans, siamang and macaque genomes, HSat1B showed fewer occurrences, and was only present in its truncated forms. H1Bsv1 showed the highest number of occurrences in the orangutans (chr. 2, 10, 17 and 23), while H1B is restricted to chr.17 and 23 and H1Bsv2 is only located on chr. 23. For siamang, H1B, H1Bsv2 and H1Bsv1 are only located on one autosome (chr. 18), except for the latter, also located on chr13. Macaque genomes showed a similar pattern with one chromosome containing all three sequences H1B and H1Bsv1 and 2 (chr. 10 in MFA and chr.15 in MNE) and again one H1Bsv1 copy also identified in another autosome (chr. 11 in MFA and chr.10 in MNE). Finally, in lemur genome we only identified a single occurrence of H1B on chr.21.

### Abundance of HSat1B monomer across *Homininae*

The chromosome-level distribution of HSat1B abundance reveals a striking pattern in humans and substantial haplotypic variation across the great apes (Figure 3, Supplementary Table S4). In the human assembly, the sequence exists mostly on chromosome Y, accounting for 92% of its total abundance, leading to the relatively small contributions from autosomes, generally varying between less than 1% and 3%.

**Figure 3.**
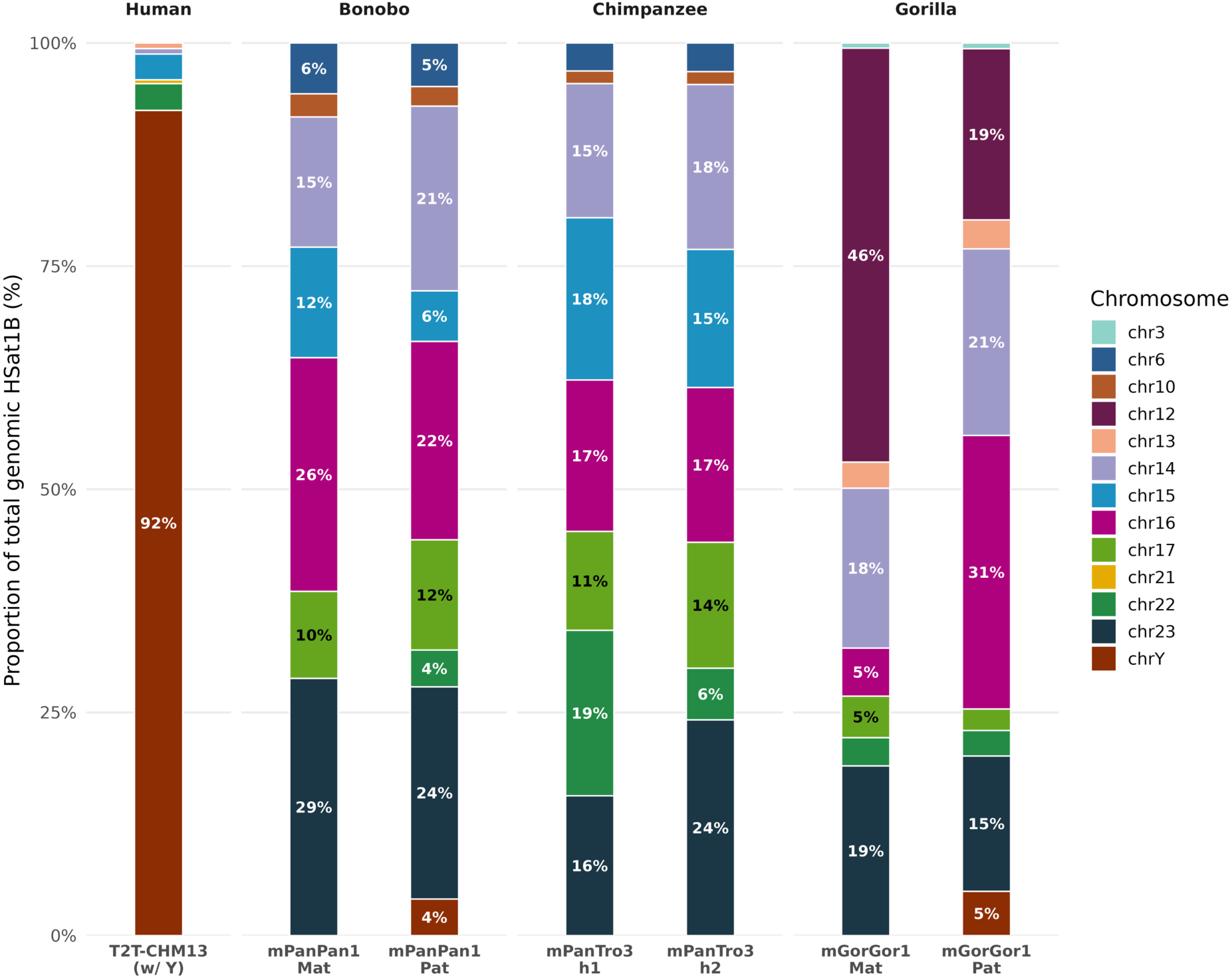
Chromosomal distribution and haplotypic variation of HSat1B monomer abundance across Homininae genomes. Stacked bar plots illustrating the relative chromosomal proportion (%) of HSat1B in human (T2T-CHM13v2.0 with Y), bonobo (mPanPan1, maternal haplotype h1 and paternal haplotype h2), chimpanzee (mPanTro3, haplotypes h1 and h2), and gorilla (mGorGor1, maternal haplotype h1 and paternal haplotype h2). Percentage labels are shown inside bars for chromosomal contributions ≥ 4% (complete base-pair counts and percentages are detailed in Supplementary Table S4).

For bonobo haplotypes, the main differences between the two haplotypes are evident on chr14 and chr15. The paternal haplotype shows a significantly larger proportion on chromosome 14 (21%) compared to the maternal haplotype (14%). Conversely, the maternal haplotype has a higher percentage on chromosome 23 (29% vs. 24% on paternal). As mentioned before, HSat1B was only identified in chr22 on the paternal haplotype corresponding to a percentage of 4%.

The chimpanzee assembly presents a higher haplotypic variation. Haplotype 2 has a higher percentage of HSat1B on chromosome 23 (24% vs. 16% in haplotype 1) and a lower percentage on chromosome 22 (6% vs. 19% in haplotype 1). On the other hand, haplotype 1 shows a lower abundance on chr14 (15% vs. 19% in haplotype 2) and chromosome 15 (18% vs. 16% in haplotype 2).

The gorilla assembly exhibits the most drastic differences between haplotypes, where HSat1B is mostly present in the maternal haplotype of chr12 (46%), which is showing a substantially higher proportion than the one in the paternal haplotype (19%), and shows a lower percentage on chromosome 16 (5% vs. 31% in paternal haplotype).

### Nuclear localization of H1B-containing RNA signals and repeat architecture of transcriptomic reads

Several satellite DNAs were shown to be transcribed into satellite non-coding RNAs, with some of them associated with relevant features in health and disease (Lopes et al., 2021). Following this reasoning, RNA-FISH was performed in human and gorilla cells using the same H1B-targeting probe employed for chromosomal mapping. Nuclear hybridization signals were detected in both species (Figure 4A), demonstrating the presence of H1B-containing RNAs. Because the probe targets the H1B region rather than the complete HSat1B monomer, these signals cannot be assigned exclusively to canonical HSat1B transcripts and may also derive from dispersed H1B-related loci.

**Figure 4.**
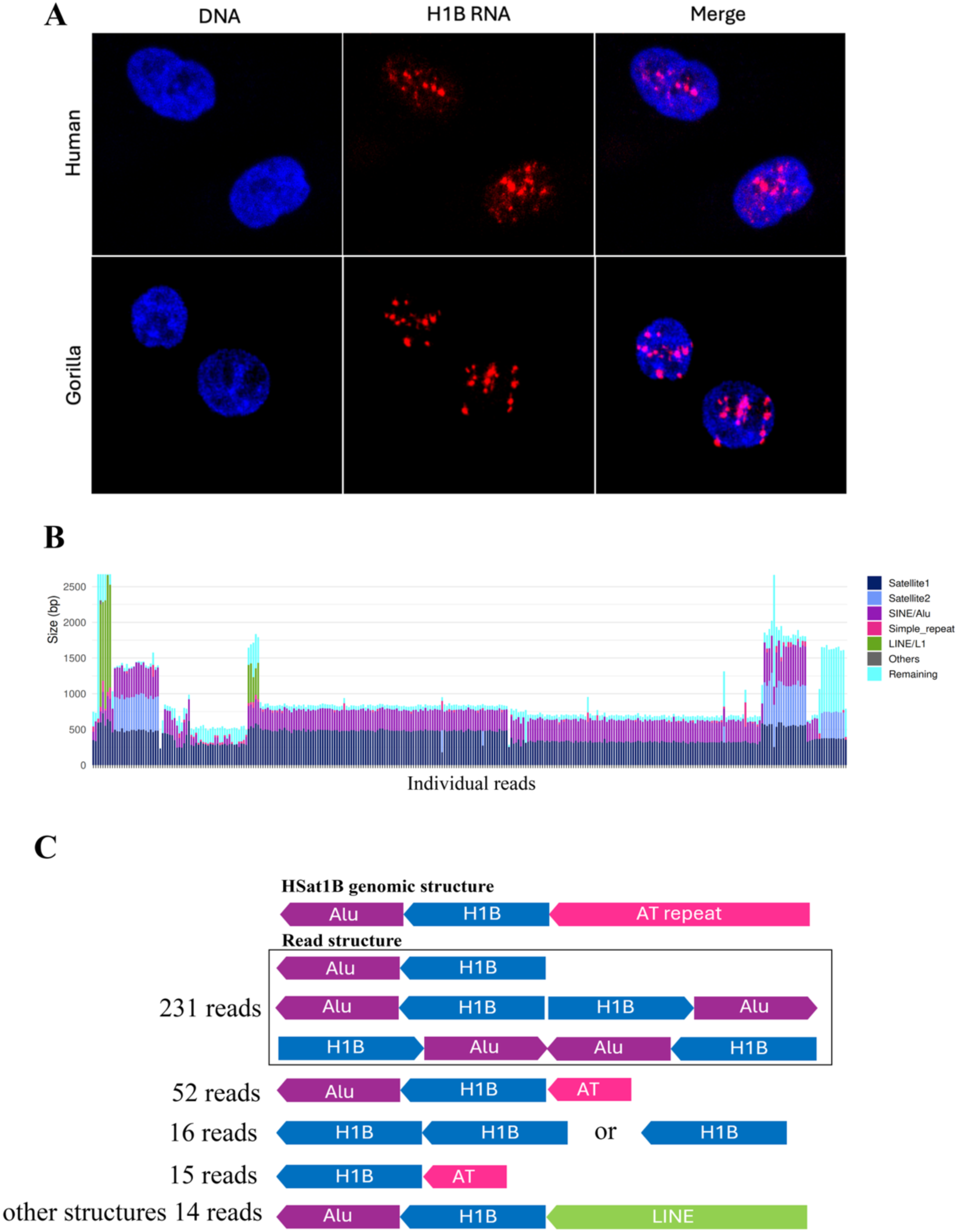
Transcriptional activity and read-level repeat architecture of H1B-containing RNAs. **(A)** Representative 3D RNA-FISH confocal microscopy images of human and gorilla interphase nuclei hybridized with the H1B-targeting probe (red) and counterstained with DAPI (blue), demonstrating the nuclear localization of H1B-containing transcripts in both species. **(B)** Read-level repeat composition of 328 H1B-containing long RNA reads extracted from a human sperm transcriptomic dataset (SRR10123831). Each vertical bar represents an individual read plotted by size (bp) and color-coded according to RepeatMasker annotation: dark blue, primary H1B (Satellite 1) segment; cyan, secondary H1B (Satellite 2) segment in reads spanning two units; purple, SINE/*Alu*-derived sequences; pink, simple repeats; green, LINE/L1-derived sequences; grey, other repeats; and light blue, remaining unannotated sequence. **(C)** Schematic comparison between the canonical HSat1B genomic monomer and the major recurrent repeat architectures identified across the 328 H1B-containing sperm transcripts, with the number of reads supporting each conformation indicated on the left.

To further understand the HSat1B non-coding transcript structure, we extracted 328 reads containing H1B sequence (Supplementary Table S5a) from a publicly available transcriptomic dataset (SRR10123831) from human sperm long-read RNA (Sun et al., 2021). The use of this dataset was since HSat1B is highly enriched on the human Y chromosome and because elevated satellite DNA transcription has been reported in the germline context (Šatović-Vukšić & Plohl, 2023). We characterized them and identified other repetitive elements in these reads, including Alu-derived sequences, simple repeats and LINE-derived elements (Supplementary Table S5b). A read-level inspection showed H1B homologous segments of up to ∼600 bp embedded within total read lengths reaching ∼3.5 kb, with a subset of reads carrying an additional adjacent satellite-annotated block of up to ∼1.1 kb (Figure 4B), illustrating the variability of satellite/TE across individual transcripts. In most reads (in 283 out of 328), satellite and Alu annotations appeared in adjacent blocks within the same read, supporting that H1B transcripts demonstrate alternative conformations (Figure 4C). The Alu sequences present in the transcripts belong to the AluS and AluY subfamilies (Supplementary Table S5c).

## Discussion

Although HSat1B remains one of the least characterized human classical satellites, our comparative analysis across ten primate species provides a detailed assessment of its structure, abundance, chromosomal distribution and evolutionary trajectory. This work shows that HSat1B is not exclusive to humans but is also present in other *Homininae* genomes, while H1B-related sequences have a broader distribution across primates.

### HSat1B is restricted to *Homininae* species

The restriction of full HSat1B arrays to *Homininae* species, together with their marked expansion on the human Y chromosome, indicates a lineage-specific amplification history. Although HSat1B is also present on the Y chromosomes of gorilla and bonobo, its much lower abundance in these species suggests that Y-linked amplification intensified in the human lineage, although assembly and sampling limitations preclude precise reconstruction of the timing and magnitude of this expansion.

The availability of phased diploid assemblies for the non-human great apes allowed the detection of substantial satellite variation between homologous chromosomes. Haplotype-specific differences were identified across all three non-human *Homininae* species and were particularly pronounced in gorilla (Figure 3), where the maternal haplotype harbored 46% of total HSat1B content on chromosome 12 compared to 19% in paternal haplotype. Such marked haplotypic variation points to ongoing evolutionary dynamism in HSat1B arrays, likely driven by unequal crossing over events that generate copy-number polymorphisms between homologous chromosomes (Altemose et al., 2022). Although the available data do not allow us to establish the precise timing of HSat1B origin or completely exclude alternative evolutionary scenarios, such as ancestral presence followed by lineage-specific loss or sequence divergence, the detection of the complete HSat1B composite monomer exclusively in the analyzed Homininae genomes strongly supports the hypothesis that this repeat organization emerged or underwent substantial expansion within the Homininae lineage (∼16 Mya) (Figure 5). This classifies HSat1B as a relatively young satellite on the primate evolutionary timescale, contrasting with ancient families such as alpha satellite, which is shared across all primates (Sujiwattanarat et al., 2015).

**Figure 5.**
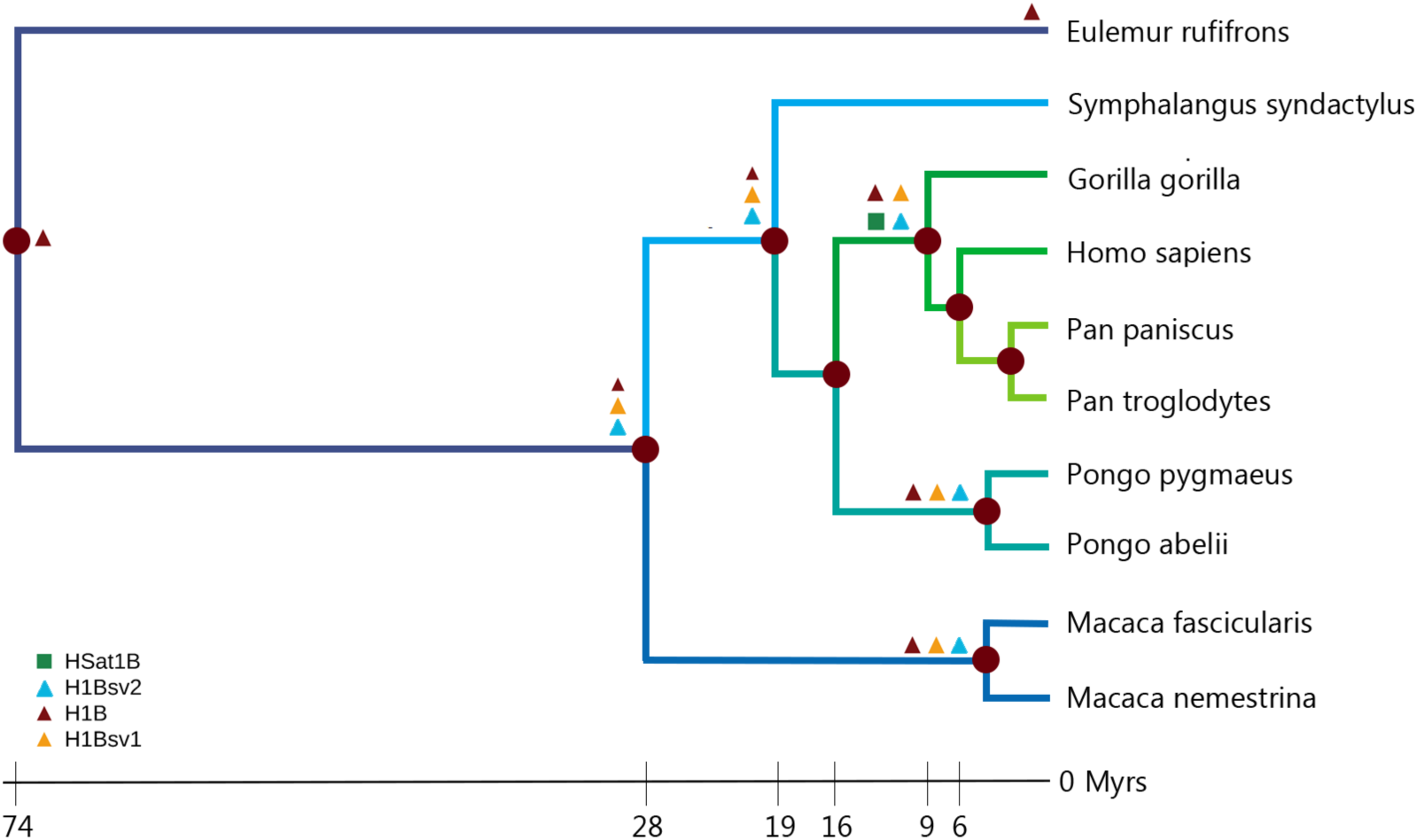
Phylogenetic reconstruction of the evolutionary emergence of H1B variants and HSat1B across primates. Primate time tree illustrates the presence and diversification of H1B (red triangles), H1Bsv1 (yellow triangles), H1Bsv2 (blue triangles), and the composite HSat1B monomer (green squares) along evolutionary branches. While dispersed H1B-related units are conserved across all analyzed primates dating back ∼74 Mya, the canonical tripartite HSat1B tandem repeat emerged specifically in the Homininae ancestor (∼16 Mya). Divergence times (in million years ago, Myrs) and tree topology are derived from TimeTree 5 (Kumar et al., 2022).

The overall agreement between DNA-FISH mapping and *in silico* predictions supports the reliability of our computational identification of major HSat1B arrays on the Y chromosome and the pericentromeric regions of acrocentric chromosomes. In contrast, the DNA-FISH signals were also detected on chromosomes 3 in both male and female cell lines, and on chromosome 4 in the male cell line, where no canonical HSat1B arrays are annotated in the T2T-CHM13v2.0 assembly (Figure 1D, 1E). Because our probe targets the H1B component rather than the full monomer, these signals may reflect dispersed H1B-related sequences, low-copy variants below the array threshold, or inter-individual structural polymorphisms not captured in the single-genome T2T-CHM13v2.0 reference. These observations underscore how sequence-based annotation and cytogenetic mapping provide complementary insights for characterizing highly repetitive genomic regions, while reinforcing the need for more diverse pangenome resources.

### HSat1B origin and dynamics is closely linked with transposable elements

The identification of H1B-related sequences across all analyzed primate genomes, including lemur, indicates that the GC-rich H1B unit predates the emergence of the complete HSat1B monomer and was likely already present in an early primate ancestor. Its occurrence as isolated copies or small clusters in TE-rich genomic contexts is consistent with a stepwise evolutionary process in which an ancestral H1B unit was subsequently incorporated into the tripartite HSat1B repeat. The broader distribution of H1Bsv1 across primate lineages may indicate that it represents an ancestral H1B-derived variant. However, its apparently broader genomic distribution should be interpreted cautiously, as structural differences between H1Bsv1 and canonical H1B, together with the similarity thresholds used for detection, may influence the number of locations detected. Furthermore, the frequent co-localization of H1Bsv1 with HSat1B arrays in great apes supports a close evolutionary link between these H1B-derived variants and the genomic regions from which composite HSat1B arrays expanded.

Our data supports a stepwise evolutionary model involving an ancestral H1B sequence and transposable elements (Figure 6A). The ancestral H1B unit was likely restricted to a single chromosomal locus in the common ancestor of primates. Subsequent segmental duplications generated multiple local H1B copies - as observed in the most phylogenetic distant species analyzed here, Bennett’s brown lemur *-* creating the local sequence redundancy required for structural innovation. (Plohl et al., 2008; Šatović-Vukšić & Plohl, 2023) The close physical association of H1B sequences with LINE and Alu elements suggests that TE activity contributed to the structural diversification of H1B into H1Bsv1 and H1Bsv2, as well as their subsequent dispersal to additional chromosomes in orangutan, macaque and siamang via TE-mediated transduction or ectopic recombination (Šatović-Vukšić & Plohl, 2023; Schmitz & Querques, 2024; Sundaram et al., 2014). Indeed, TEs are well-established drivers of sequence mobilization and genomic plasticity, and prior TE insertions can render genomic regions preferential hotspots for subsequent retrotransposition events (Palomeque et al., 2006; Schmitz & Querques, 2024). H1Bsv2 arose through the insertion of an Alu element within H1B and was maintained along primates’ evolution. Previous works suggest that Alu insertions preferentially occur in relatively GC-rich genomic regions (Lee et al., 2008), making H1B containing regions preferable insertion sites given their high content in GC. Moreover, H1Bsv2 was consistently found on a single chromosome, in proximity to H1B and H1Bsv1 sequences across the analyzed species and close to HSat1B arrays in *Homininae* species. This pattern may indicate an ancestral translocation event that was inherited across primate lineages or, less likely, independent Alu insertions at a shared genomic hotspot (Palomeque et al., 2006; Schmitz & Querques, 2024).

**Figure 6.**
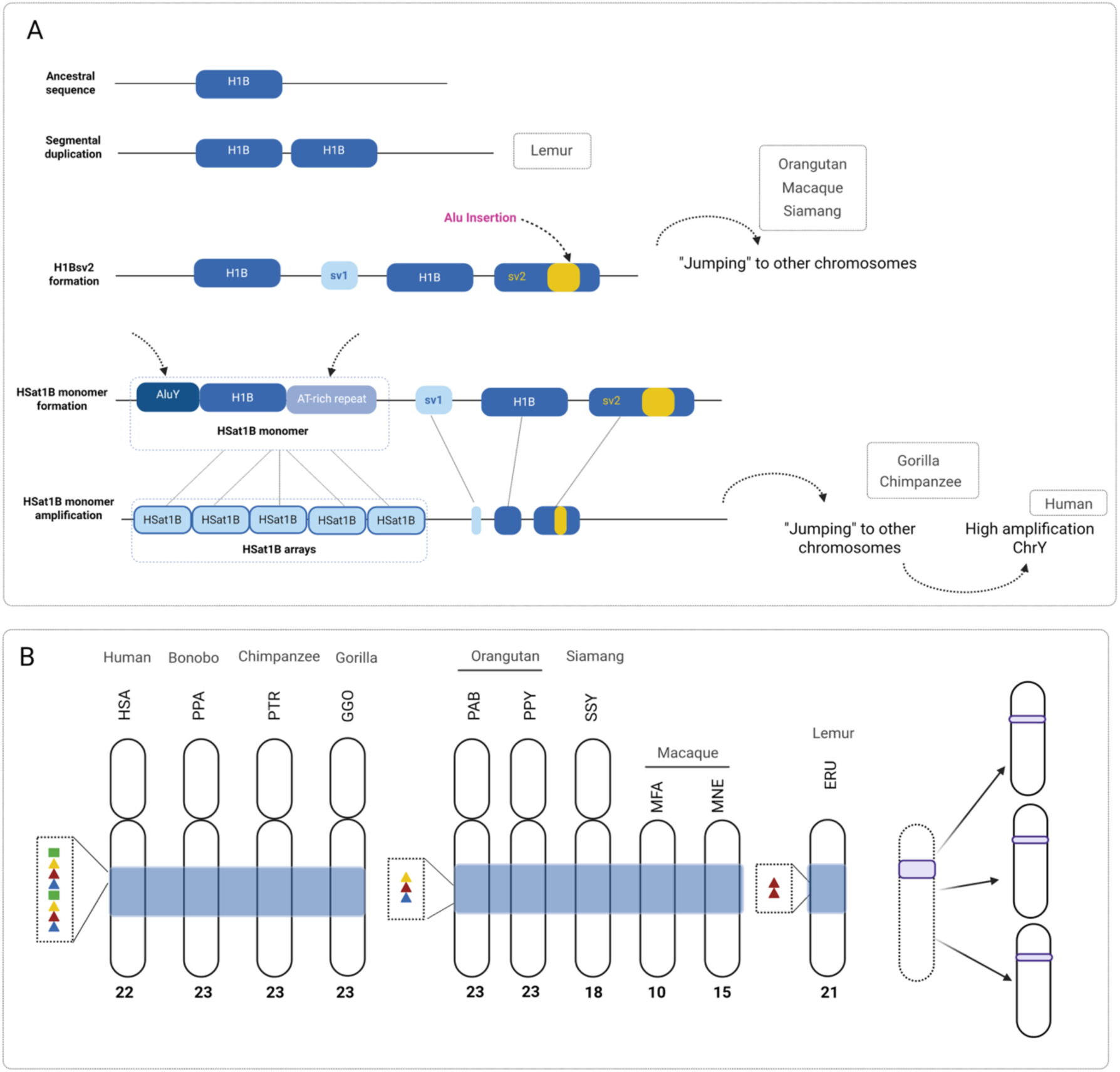
Proposed evolutionary model of HSat1B birth, diversification, and chromosomal dispersal across primates. **(A)** Stepwise model for the origin and amplification of HSat1B. An ancestral single-copy H1B unit present in the primate ancestor underwent local segmental duplication (as retained in lemur). Subsequent sequence divergence and transposable element activity generated the structural variants H1Bsv1 and H1Bsv2 - the latter via an *Alu* insertion within the H1B unit (pink arrow and yellow block) - followed by inter-chromosomal dispersal in Old World monkeys, lesser apes, and orangutans. In the Homininae ancestor, the juxtaposition of an *AluY* element and an AT-rich repeat block adjacent to H1B formed the composite ∼2.5 kb HSat1B monomer. Tandem amplification generated canonical HSat1B arrays, which subsequently spread across chromosomes in gorilla, bonobo, and chimpanzee, and underwent massive lineage-specific amplification on the human Y chromosome. **(B)** Comparative synteny ideogram of orthologous primate chromosomes harboring the ancestral H1B/HSat1B cluster (shaded blue region): human (HSA, chr22), bonobo (PPA, chr23), chimpanzee (PTR, chr23), gorilla (GGO, chr23), Bornean orangutan (PAB, chr23), Sumatran orangutan (PPY, chr23), siamang (SSY, chr18), crab-eating macaque (MFA, chr10), pig-tailed macaque (MNE, chr15), and Bennett’s brown lemur (ERU, chr21). The schematic on the right illustrates how a single ancestral locus (purple band) diversified and dispersed onto additional chromosomes during primate evolution, conceptually summarizing the lineage-specific patterns shown across the ideograms. Created with BioRender.com.

The insertion of an AluY-derived element adjacent to H1B, together with the expansion of an AT-rich simple repeat, likely contributed to the formation of the complete HSat1B monomer. Subsequent amplification of this composite unit would have enabled the generation of the extended HSat1B arrays observed in *Homininae* genomes.

The spread of HSat1B and related sequences across the human acrocentric chromosomes (13, 14, 15, 21, and 22) is readily explained by their physical clustering within the nucleolus around Nucleolar Organizer Regions (NORs) (Bersaglieri & Santoro, 2019), which facilitates inter-chromosomal sequence exchange between non-homologous chromosomes. Moreover, recent T2T pangenome analyses have demonstrated that the short arms of human acrocentric chromosomes share pseudo-homologous regions (PHRs) that undergo frequent heterologous recombination (Guarracino et al., 2023), homogenizing satellite arrays across acrocentric pairs. The conservation of H1B- and HSat1B-related sequences within a syntenic region corresponding to human chromosome 22 across the analyzed primate genomes may represent an ancestral genomic context from which related sequences were subsequently dispersed. The close physical association of acrocentric chromosomes within nucleoli may have facilitated subsequent interchromosomal exchange of these sequences. However, this mechanism alone does not account for the markedly larger HSat1B content observed on the human Y chromosome, which likely reflects an additional, lineage-specific amplification event. Similarly, the apparent absence of HSat1B from the chimpanzee Y chromosome, despite its presence on the bonobo Y chromosome, may reflect either a lineage-specific loss, differential array retention, or limitations in the assembly or annotation of this highly repetitive region. This is directly supported by our H1B clustering analysis, in which monomer sequences from chromosomes 13, 21 and 22 frequently collapsed into shared 99%-identity clusters, providing molecular evidence consistent with sequence exchange between acrocentric pseudo-homologous regions (Supplementary Figure 1). In the human genome, HSat1B underwent massive amplification on the Y chromosome after the divergence of the human lineage. However, the detection of HSat1B on the Y chromosomes of gorilla and bonobo indicates that the association of this satellite with the Y chromosome predates the divergence of these *Homininae* lineages. Therefore, the ancestral event was likely the recruitment or initial establishment of HSat1B on the Y chromosome, whereas the extensive Y-linked expansion observed in humans represents a later, human-specific amplification. This is also supported by recently published data, reporting the occurrence of recent structural rearrangements involving a 5Mb region on human Y chromosome, spanning blocks of HSat1B and HSat3, consistent with evolution mechanisms of unequal exchange followed by sequence homogenization (Rhie et al., 2023).

Growing evidence shows that satellite DNAs and transposable elements are deeply interconnected, with TEs often facilitating the dispersal and diversification of satDNA repeats (Šatović-Vukšić & Plohl, 2023). Scalvenzi and Pollet (Scalvenzi & Pollet, 2014) proposed that transposable elements can capture and internally amplify satellite DNA, dispersing it across the genome to seed traditional satellite arrays that expand through non-reciprocal recombination. More recently, Šatović-Vukšić et al. showed that satDNA formation can involve tandemization of DNA segments, often facilitated by TE-associated sequence motifs, generating novel repeats that retain short regions of sequence homology to TEs within their monomer units. Our data is consistent with this TE-mediated model. The presence of an AluY-derived fragment within the HSat1B monomer, the occurrence of H1B-related sequences in TE-rich genomic contexts, and the Alu insertion characterizing H1Bsv2 suggest that Alu elements may have contributed to the structural diversification and subsequent incorporation of H1B-derived sequence into the HSat1B repeat unit. Furthermore, the detection of AluS and AluY sequences, as well as LINE-derived segments, within H1B-containing sperm transcripts supports the idea that these satellite-derived transcripts are embedded in a broader repeat-rich genomic and transcriptional environment. Together, these observations support a stepwise model in which TE activity contributed to H1B diversification, HSat1B monomer formation and the subsequent amplification of HSat1B arrays.

### HSat1B is transcriptionally active in primates

Transcription of satellite DNA into satellite non-coding RNAs(satncRNAs) is increasingly recognized across eukaryotic genomes. For instances, human Satellite 1A (HSat1A) is transcribed by RNA polymerase II into polyadenylated lncRNAs whose expression varies with cell type and physiological state (Lopes et al., 2021, 2023). Here, RNA-FISH analysis revealed distinct nuclear signals for H1B-containing RNAs in both human and gorilla cell lines (Figure 4**A**), demonstrating that this sequence module is transcriptionally active across Homininae species. Because our probe targets the core H1B unit rather than the full tripartite monomer, the relative transcriptional contribution of canonical HSat1B arrays versus dispersed H1B-related loci remains to be determined. Nevertheless, analogous to the roles of HSat3 transcripts in the cellular stress response (Giordano et al., 2020) and HSat1A transcripts in pericentromeric regulation (Lopes et al., 2023), H1B-containing transcripts may participate in heterochromatin maintenance, subnuclear architecture, or chromosome-specific regulatory pathways. Notably, pericentromeric satellite RNAs often facilitate the recruitment of heterochromatin proteins such as HP1 and histone methyltransferases, and the GC-rich H1B unit has been proposed to support specialized chromatin conformations required for pericentromeric heterochromatin assembly (Ninomiya et al., 2023).

The repeat composition of H1B-containing sperm transcripts suggests that these RNAs originate from, or are processed through, genomic regions enriched in diverse repetitive elements. Their variable content, including Alu-derived, LINE-derived and simple repeat sequences, is consistent with transcription from heterogeneous H1B/HSat1B-related genomic contexts rather than from a single canonical monomer. This interpretation is supported by the recurrent proximity of H1B-related genomic sequences to Alu and LINE-1 elements, as well as by the known capacity of TEs to influence the transcriptional output of neighbouring genomic regions (Šatović-Vukšić & Plohl, 2023; Schmitz & Querques, 2024). The LINE-derived sequences detected in these transcripts most likely reflect LINE-1 elements located within or near the genomic regions from which the H1B-containing RNAs are derived, rather than indicating that H1B itself is LINE-derived. Their presence is nevertheless relevant because LINE-1 activity provides the reverse transcriptase required for the retrotransposition of non-autonomous Alu elements (Dewannieux et al., 2003). Alu elements remain active in the human genome, where de novo insertions continue to occur, although their retrotransposition rate has varied over evolutionary time (Cordaux et al., 2006) resulting in Alu elements present in introns, protein-coding exons, as well as non-coding regions (Häsler et al., 2007). Although Alu elements were initially assumed to be transcriptionally silent, the identification of RNA polymerase II and III-derived Alu transcripts confirmed their expression as independent non-coding RNAs (Gussakovsky & McKenna, 2021). These standalone Alu RNAs remain less characterized than Alu sequences embedded within mRNAs or introns. The sequence differences between H1B-containing transcripts and their genomic representations may reflect transcription from diverse H1B-related loci, post-transcriptional processing, or incomplete representation of repetitive arrays in the reference assembly. Although these differences can complicate the unambiguous assignment of individual transcripts to a single chromosomal origin, particularly within large and highly homogeneous satellite arrays, the specific combination of H1B, Alu and other repeat-derived sequences may help prioritize candidate source loci. The detection of both AluS and AluY sequences in the transcripts, whereas canonical HSat1B monomers contain an AluY-derived fragment, further suggests that some transcripts may derive from H1B-related genomic contexts other than canonical HSat1B arrays. Given that HSat1B monomers incorporate an Alu-derived segment and that H1B-containing transcripts were detected by RNA-FISH in both human and gorilla cells, the presence of repeat-associated RNAs in sperm may provide an additional framework in which H1B-derived or HSat1B-like transcripts can be considered part of a broader repertoire of repetitive RNAs associated with the primate germline. However, because H1B/HSat1B-related transcription has so far been assessed only in adherent human and gorilla cell lines and in sperm-derived transcriptomic data, its expression in additional cell types, developmental stages or physiological conditions remains to be investigated.

## Materials and methods

### Genome assemblies and data sources

Genomic analysis used publicly available T2T and chromosome-scale assemblies for 10 primate species: *Homo sapiens* (T2TCHM13v2.0), *Pan paniscus* (mPanPan1), *Pan troglodytes* (mPanTro3), *Gorilla gorilla* (mGorGor1), *Pongo pygmaeus* (mPonPyg2), *Pongo abelii* (mPonAbe1), *Symphalangus syndactylus* (mSymSyn1), *Macaca nemestrina* (mMacNem1), *Macaca fascicularis* (T2T-MFA8) and *Eulemur rufifrons* (OSU_ERuf_1). Assemblies and annotations were obtained from the UCSC Genome Browser and NCBI databases (full list of GenBank and RefSeq assembly accessions on Supplementary Table S2).

### Identification and annotation of HSat1B and H1B sequences

RepeatMasker (v4.2.1;Nishimura, 2000) with FamDB v2.0.5 and RepBase libraries identified HSATI/HSat1B regions in each assembly. A representative HSat1B monomer sequence (Clone 3) was used via the NCBI BLAST web server (Altschul et al., 1990) query against all primate genomes to detect array regions and dispersed H1B-related sequences (word size = 7, E-value ≤ 1×10^-5^). These parameters were chosen to balance sensitivity to divergent H1B-related sequences with specificity against false short matches. H1Bsv1 and H1Bsv2 were identified through manual inspection of BLAST output alignments. Coordinates for HSat1B, H1B, H1Bsv1 and H1Bsv2 loci in non-human primates are provided in Supplementary Tables S3a–i. Coordinates for human H1B and variant locations are provided in Supplementary Table S1b. H1B sequences detected by RepeatMasker were aligned using the MAFFT web server and the dotplot generated by the platform was used to compare each candidate against the canonical H1B and confirmed sequence variants (dotplots available on additional data). Hits lacking the expected alignment pattern were discarded and are available on additional data. To further characterize transcript-associated repeat context, the H1B sequence (chrY:35018931-35019496) was used as an additional BLAST query and the corresponding hit sequences were retrieved as FASTA. Sequences with less than 40% query coverage were excluded and the retained sequences were reannotated with RepeatMasker. Retained hit sequences were additionally checked via pairwise dotplot comparison using the MAFFT web server for confirmation of repeat structure prior to RepeatMasker reannotation. Representative dotplots confirming H1B repeat structure in retained transcript-associated reads are provided in the Additional Data. Transcriptome sequences were obtained from a published human sperm long-read RNA dataset (Sun et al., 2021) by extracting reads bearing HSATI/H1B-related RepeatMasker annotations and applying the same H1B-based BLAST and coverage filters used for genomic sequences. HSat1B arrays were defined as clusters of at least 2 monomers and pairwise identity ≥ 80%, whereas isolated H1B hits outside such clusters were classified as dispersed H1B. The genomic location of HSat1B, H1B, H1Bsv1 and H1Bsv2 loci was assessed by intersecting their coordinates with RepeatMasker TE annotations using BEDTools (v2.x). HSat1B and H1B abundances per chromosome were calculated by summing total base pairs covered by BLAST hits above defined identity and length thresholds. To examine the repeat environment of HSat1B- and H1B-associated sequences in genome assemblies, RepeatMasker annotations were parsed for regions overlapping HSat1B/H1B-derived hits, repeat classes and families were summarized and representative read-level annotations were selected for visualization based on the presence of adjacent HSATI and Alu-derived segments.

### *In silico* chromosomal mapping and synteny analysis

Chromosomal distributions were visualized using the RIdeogram package in R. Synteny analysis compared to the chromosomal locations of HSat1B and H1B across species using the T2T-CHM13v2.0 human genome assembly as reference. Syntenic relationships were assessed in the NCBI Multiple Comparative Genome Viewer by comparing local gene order and centromere-proximal positions across orthologous chromosomes.

For Figure 3, chromosome-level HSat1B abundance was expressed as the percentage contribution of each chromosome to the total HSat1B content within each assembly or haplotype. These proportions were visualized as stacked bar plots using ggplot2 in R. Chromosomes were ordered consistently across species, and percentage labels were displayed for chromosomal contributions of at least 4%.

### Sequence clustering and phylogenetic analysis of H1B monomers

H1B monomer sequences extracted from the human (T2TCHM13v2.0), gorilla (mGorGor1, maternal and paternal haplotypes), bonobo (mPanPan1, haplotypes 1 and 2) and chimpanzee (mPanTro3, haplotypes 1 and 2) assemblies were clustered using CD-HIT-EST (99% identity threshold, word size 10) to identify near-identical sequence groups and remove redundancy prior to phylogenetic inference. Clustering was not applied to bonobo and chimpanzee datasets given their comparatively low sequence counts. Clustered representative sequences were aligned using MAFFT (--auto) and maximum-likelihood phylogenetic trees were inferred using IQ-TREE (ModelFinder Plus for model selection, 1000 ultrafast bootstrap replicates). The full workflow was implemented in Snakemake. Cluster membership was additionally used to assess sequence identity sharing between chromosomal locations: clusters containing monomers from more than one chromosome were classified as cross-chromosomal.

### Cell culture, DNA-FISH and RNA-FISH

Human lymphoblastoid GM12878 (female, XX) and fibroblast HDFn (male, XY) cell lines, as well as male gorilla (*Gorilla gorilla*) fibroblasts, were cultured under standard conditions. Lymphoblastoid cells were maintained in RPMI 1640 medium supplemented with 10% fetal bovine serum and antibiotics at 37 °C in a humidified 5% CO2 incubator. Fibroblasts were grown in Advanced DMEM supplemented with AmnioMAX C100 and fetal bovine serum under the same conditions. Chromosome spreads were obtained by adding colcemide (0.1 µg/mL) for 1 h to arrest cells in metaphase, followed by hypotonic treatment with 0.075 M KCl and fixation in methanol:acetic acid (3:1, v/v). An HSat1B probe was made by PCR labeling with biotin-16-dUTP (Roche), using HSat1B-specific primers (Primer Forward HSat1B – 5’-GTCTTTCAAAGGTCAGAAGA-3’; Primer Reverse HSat1B – 5’-CATAACCGATGAAACCTACT-3’) on human genomic DNA. The PCR product was cloned into a plasmid vector; colonies were selected by blue-white screening and inserts were verified by Sanger sequencing. For DNA-FISH, metaphase slides were baked at 65°C for 1 hour and denatured in 70% Formamide solution at 65°C for 1min and 30 sec. Probe was denatured at 65°C for 15 minutes and hybridization was performed over-night. Post-hybridization stringency was done with 1×SSC at 73°C. Biotin-labelled H1B probe was detected with Streptavidin Cy3 conjugated (Sigma-Aldrich). Preparations were mounted with mounting media containing 4-6-diamidino-2-phenylindole (DAPI) (Vector Laboratories) to counterstain chromosomes. DNA-FISH images were observed using a Zeiss ImagerZ2 microscope coupled to an ORCA-Flash 4.0 digital camera (Hamamatsu) and captured using SmartCapture 4 software (Digital Scientific, UK).

RNA-FISH was performed following Ferreira et al. (2020), with some modifications. Cells were hybridized overnight at 37°C with the Biotin-labelled H1B probe. Probe detection was carried out with Streptavidin, Cy3 conjugated, (Sigma-Aldrich). Cells were mounted with coverslips and counterstained with mounting medium containing 4-6-diamidino-2-phenylindole (DAPI) (Vector Laboratories). Confocal fluorescence imaging was performed on an LSM 510 META with a Zeiss Axio Imager Z1 microscope and LSM 510 software (version 4.0 SP2). Applied settings were constant. Used lasers were helium neon (543 nm) set at 50.8%, and diode (405 nm) set at 9.9%. The pinhole was set to 102 µm (0.98 Airy units) for helium neon laser, and 112 μm for the diode laser using a 63 × objective. Images were captured at a scan speed of 5 with 1 µm thick Z-sections. AutoQuant X3 software (Media Cybernetics) allowed 3D deconvolution. Subsequent TIFF processing was run in ImageJ (1.52v). The three-dimensional isosurfaces and orthogonal slices (perpendicular or parallel angles) were produced in Image Pro Premier 3D (version 10, Media Cybernetics).

## Code availability

The R script used for ideogram generation, together with related plotting scripts, is available on Additional Data.

## Ethics statement

All cell lines used were obtained from established repositories under their existing ethical approvals; no new animal or human sampling was performed for this study

## Author contributions

**Carlos Vila-Verde**: Conceptualization, Methodology, Software, Formal analysis, Investigation, Data curation, Visualization, Writing – original draft, Writing – review and editing. **Alessia Daponte**: Investigation, Formal analysis, Validation, Visualization, Writing – review and editing. **Daniel Eleutério**: Investigation, Formal analysis, Validation, Visualization, Writing - review and editing. **Raquel Chaves**: Formal analysis, Writing – review and editing. **Claudia Rita Catacchio**: Methodology, Resources, Supervision, Writing – review and editing. **Sandra Louzada**: Conceptualization, Methodology, Resources, Supervision, Project administration, Writing – review and editing.

## Acknowledgments

This work was supported by the Scientific Employment Stimulus contract CEECIND/01825/2017 (10.54499/CEECIND/01825/2017/CP1 418/CT0003) (to SL). We also acknowledge financial support under the National Recovery and Resilience Plan (NRRP),Mission 4,Component 2,Investment 1.1, Call for tender No. 104 published on 2.2.2022 by the Italian Ministry of University and Research (MUR), funded by the European Union – Next Generation EU – Project Title “Insights into the fast genome evolution of Gibbons through single-cell strand sequencing and simulation-based approaches” – CUP H53D23001720006 - Grant Assignment Decree No. 970 adopted on 30/06/2023 by the Italian Ministry of University and Research (MUR) (C.R.C.).

---

**Supplementary Figure 1.**
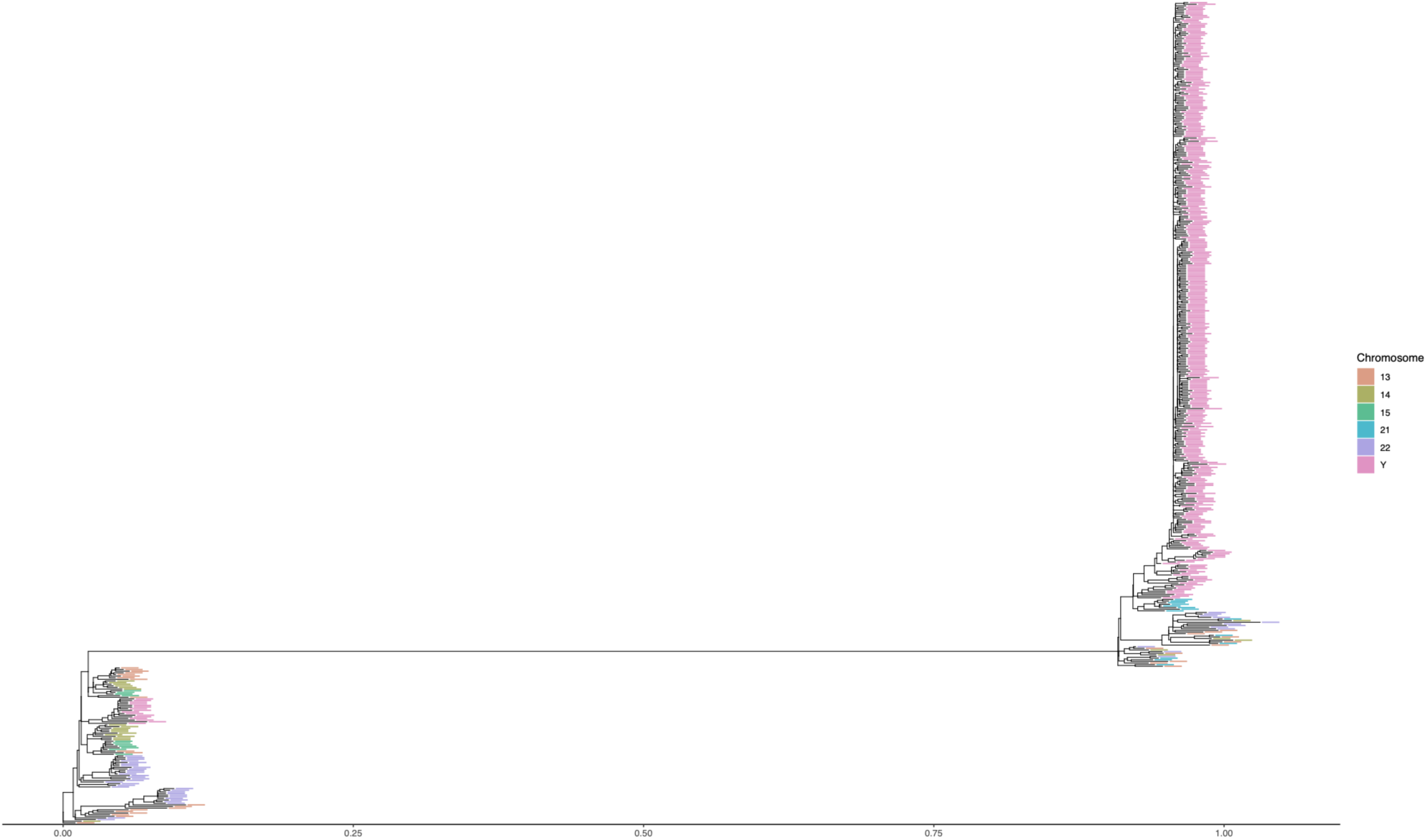
Phylogenetic relationships among human H1B sequences. Maximum-likelihood phylogenetic tree inferred from H1B sequences identified in the human T2T-CHM13v2.0 assembly. Terminal branches are coloured according to the chromosome of origin: chromosome 13 (salmon), chromosome 14 (light green), chromosome 15 (green), chromosome 21 (cyan), chromosome 22 (blue) and chromosome Y (pink). Most chromosome Y-derived H1B sequences cluster in a distinct, densely represented group, consistent with extensive recent expansion and sequence homogenization of H1B-containing HSat1B arrays on the human Y chromosome. In contrast, H1B sequences from acrocentric chromosomes are distributed across multiple mixed-chromosome clades, consistent with sequence sharing or inter-chromosomal exchange among acrocentric regions. The scale bar represents nucleotide substitutions per site.

